# Serum from patients with Idiopathic inflammatory myopathy induces skeletal muscle weakness

**DOI:** 10.1101/2024.04.02.587168

**Authors:** Cecilia Leijding, Kristofer M. Andreasson, Begum Horuluoglu, Angeles Shunashy Galindo-Feria, Eveline Van Gompel, Maryam Dastmalchi, Stefano Gastaldello, Helene Alexanderson, Ingrid E. Lundberg, Daniel C. Andersson

## Abstract

Idiopathic inflammatory myopathies (IIM) are a group of systemic autoimmune inflammatory disorders that primarily affect striated muscles leading to weakness and accelerated fatigue. The disseminated muscle phenotype points to systemic humoral factors as mediators of the disease. Autoantibodies are important biomarkers for disease classification; it is not known if they play a direct role in IIM disease development. This study aims to investigate if IIM patient serum or isolated IgG could directly impair contractile function in muscle.

Isolated flexor digitorum brevis (FDB) muscles from healthy mice were exposed to serum (10-50%) from healthy controls or patients with recent onset IIM. Some muscles were exposed to isolated total IgG (50 or 150μg/ml) from patients with IIM. Muscle force in whole muscles was measured before and after exposure to sera or IgG. Muscle force and intracellular [Ca^2+^] in single muscle fibers were measured after exposure to serum.

FDB muscles exposed to serum from patients with IIM displayed a marked reduction in force production in both 10% and 50% serum. Moreover, single myofibers dissected from FDB muscles exposed to IIM sera displayed lower force but unaffected Ca^2+^ release during contractions, which indicates myofibrillar dysfunction, but not intracellular Ca^2+^ release, as the cause to weakness. FDB muscles exposed to total IgG from IIM patients did not display any reduction in muscle force.

In conclusion IIM patient serum, but not total IgG, impairs force production in skeletal muscle fibers. As the experiments were performed in isolated muscle, our results cannot be explained by infiltrating immune cells, impaired neuronal or vascular functions. This suggests that humoral factors play a direct role in the pathogenesis of muscle weakness in recent onset IIM.

## Letter

Idiopathic inflammatory myopathies (IIM) are a group of systemic autoimmune inflammatory disorders characterized by symmetrical skeletal muscle weakness and accelerated fatigue also in the early non-atrophic disease stage^1^. Inflammatory cell infiltrates in the muscles are commonly present and important for diagnosing IIM. Yet, the degree of infiltrates does not correlate well with muscle weakness^2^ pointing to systemic factors as important mediators of muscle weakness in IIM^3^. IgG autoantibodies are present in ∼80% of IIM patients and are important biomarkers used for subclassification and prognosis. It is debated if autoantibodies are mere biomarkers or play a causative role in disease development of IIM^4^. Whether autoantibodies target the muscle (e.g. circulating IgG targeting antigens within the muscle) causing weakness in IIM is currently an open scientific question.

In this study, we investigated if sera or isolated IgG from patients with IIM may cause muscle weakness by directly impairing contractile function of muscle fibres.

To test the hypothesis, we developed a humanized *ex vivo* experimental platform where muscles from healthy mice were exposed to serum from patients with recent onset IIM^5^. Isolated flexor digitorum brevis (FDB) muscles from healthy mice were exposed to sera (10-50% sera diluted in physiological solution) from healthy controls or patients with IIM. In some experiments, muscles were exposed to isolated total IgG from patients with IIM. Measurements of force in whole muscles were performed before and 24h after incubation in serum or IgG. Muscle force and intracellular [Ca^2+^] were measured in single FDB muscle fibers after 24h of incubation.

Exposure to sera from healthy controls did not induce significant reduction of muscle force across stimulation frequencies at 10% serum concentration, which established that mouse muscles can viably be incubated with human serum for 24h (Figure 1A). In contrast, serum from patients with IIM induced a marked reduction in force production across stimulation frequencies in both 10% and 50% serum concentration (Figure 1A). This suggests that serum from patients with IIM can induce muscle weakness. Reduced muscle force can in principle be due to reduced number of contracting myofibers and/or reduced contractile function within each myofiber. The latter may be due to impaired myofiber [Ca^2+^] release and/or impaired function at the contractile myofilaments, e.g. reduced Ca^2+^ sensitivity^6^. To investigate this, we measured tetanic force and [Ca^2+^] in intact mechanically isolated single myofibers from serum exposed FDB muscles. Tetanic force in single myofibers was lower when exposed to sera from patient with IIM compared to the controls, while tetanic [Ca^2+^] during contractions was unaffected by the IIM sera (Figure 1B). This indicates that muscle weakness is due to impaired myofibrillar function rather than perturbed intracellular [Ca^2+^] release.

**Figure 1.**
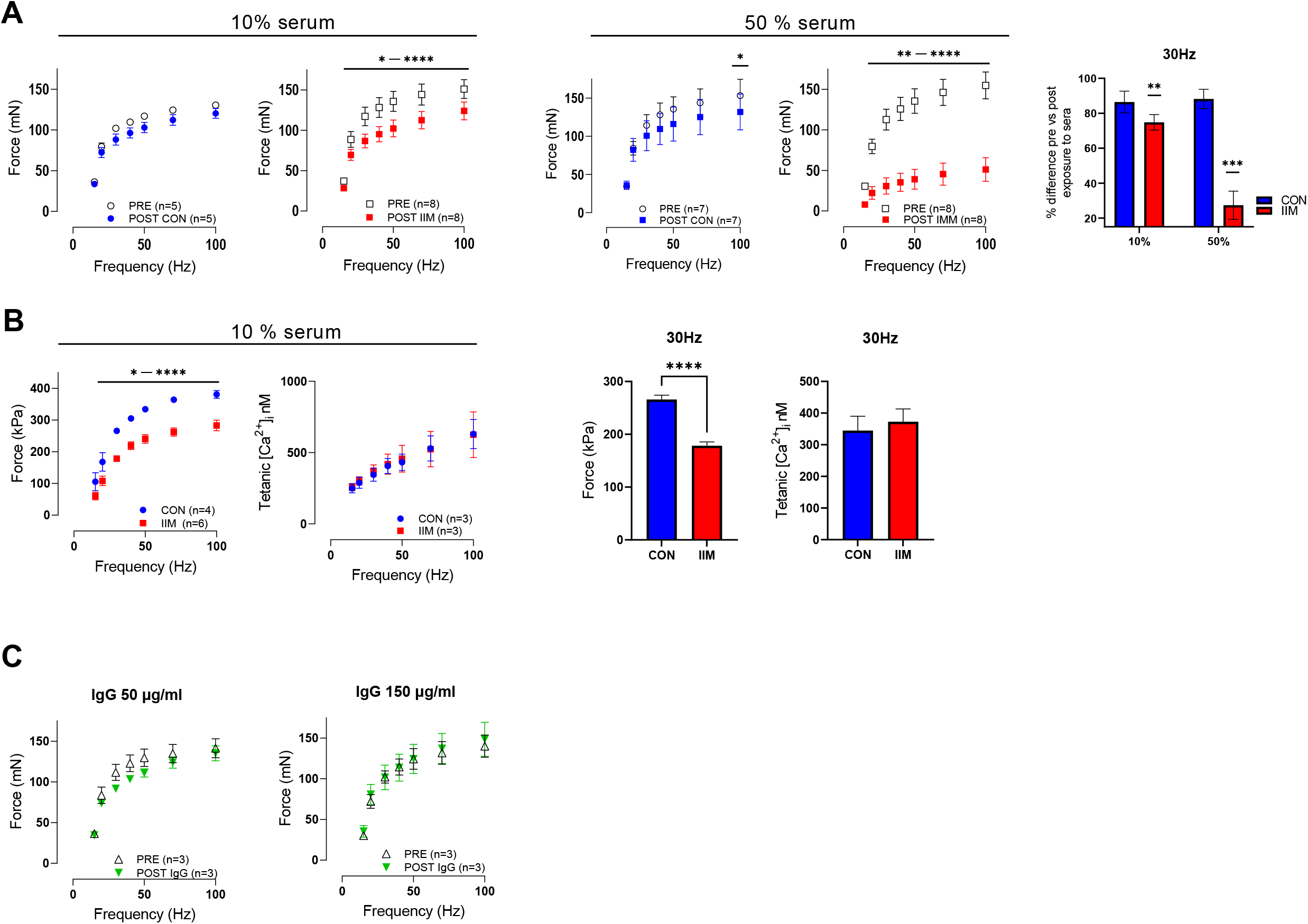
Serum from human patients with Idiopathic inflammatory myopathy (IIM) reduces contractile force in isolated skeletal muscles from healthy mice. A) Force-frequency curves from flexor digitorum brevis (FDB) muscles before (PRE) and after (POST) exposure to 10% and 50% serum from healthy control (CON; blue) or patients with idiopathic inflammatory myopathies (IIM; red). Data presented as mean ± SEM, * P < 0.05, *** P < 0.001, **** P < 0.0001, two-way ANOVA, Sidak’s multiple comparisons test performed for force-frequency measurements and paired t-test performed for 30Hz. B) Force-frequency and tetanic [Ca^2+^]_i_ -frequency curves from mechanically dissected single fibers from FDB muscles exposed to 10% serum from healthy controls (CON;blue) and IIM (IIM;red). Data presented as mean ± SEM, * P < 0.05, **** P < 0.0001, Mixed-effects analysis, Sidak’s multiple comparisons test performed for force-frequency measurements and unpaired t-test performed for force and intracellular tetanic Ca^2+^ ([Ca^2+^]_i_) at 30Hz. C) Force-frequency curves from FDB muscles before (PRE) and after (POST) exposure to total IgG from patients with IIM (50 or 150μg/ml IgG).

To test if the muscle weakness induced by IIM serum was caused by autoantibodies, we exposed muscles to isolated total IgG from patients with IIM. Interestingly, isolated total IgG did not cause any reduction in muscle force, which argues against that IgG from patients with IIM in itself could induce muscle weakness (Figure 1C).

For the first time, we have demonstrated that serum from patients with IIM, but not isolated IgG, may induce weakness directly in skeletal muscle fibers. The experiments are performed in isolated muscles, thus our results cannot be explained by infiltrating immune cells, neuronal or vascular functions. This indicates that non-cellular factors within the systemic circulation are part of the pathogenic cues that can impair muscle function in recent onset IIM.

## Contributors

CL, DCA, IEL contributed to design of the study. ASGF, BH, CL, DCA, EVG, HA, IEL, KMA, MD, SG were involved in the acquisition of data. CL, DCA, IEL, contributed to the analysis and interpretation of data. All authors contributed to drafting and/or revising the manuscript.

## Ethics approval

This study involves human samples approved by the Stockholm Ethics Examination Authority (2016/2444-31; 2018/1350-32; 2005/792-31/4) and tissue from murine approved by laboratory animal ethics committee at the Swedish Board of Agriculture (2155-2020).

## Supplemental material

Methodological description

