## Supplemental material for "Serum from patients with Idiopathic inflammatory myopathy induces skeletal muscle weakness"

Methodological description

***Experimental setting***

Intact flexor digitorum brevis (FDB) muscles were dissected from female WT (C57BL/6JRj) mice, and thereafter incubated in physiological buffer solution (Dulbecco´s Modified Eagle´s Medium/F-12 (DMEM/F-12)) containing either 10% or 50% of serum from patients with Idiopathic Inflammatory Myopathies (IIM) or from healthy controls. Serum from 4 patients and 2 healthy controls were used for experiments in 10% serum, repeated 1-3 times in the different muscles. Serum from 3 patients and 2 healthy controls were used for experiments in 50% serum, repeated >2 times in different muscles. Altogether, serum from 5 patients and 2 healthy controls were used in the experiments. In some experiments, total IgG obtained from patients with IIM (n=3) were diluted in DMEM/F-12 (50ug/ml or 150ug/ml total IgG) and thereafter incubated with FDB muscles. The isolated muscles were incubated at room temperature for ~24h.

***Force measurements whole muscles***

Force was measured in isolated FDB muscles, paired before-after, incubation with serum or IgG.

First the FDB muscle was dissected and the proximal tendon and three medial toe tendons of the muscle were tied with surgical suture thread and mounted to hooks connected to a force transducer in a stimulation chamber containing Tyrode solution ((in mM): 121 NaCl, 5.0 KCl, 1.8 $CaCl_2$, 0.5 $MgCl_2$, 0.4 $NaH_2PO_4$, 24.0 $NaHCO_3$, 0.1 EDTA, and 5.5 glucose). The Tyrode was constantly bubbled with 95% $O_2$ - 5% $CO_2$ and temperature was set to 31°C. When mounted, the muscle was stretched until maximal force-length relationship was reached. Force was assessed at increasing the stimulation frequencies from 15Hz to 100Hz, each for a duration of 350ms. The muscle had >1 min of rest between every stimulation step to avoid fatigue to develop. Contractility was measured by the FDB muscle's absolute force (mN) production for all groups.

***Isolation of total IgG***

Affinity chromatography was performed to purify IgG autoantibodies from serum as previously described [1 2]. Briefly, serum samples were centrifuged at 2000g for 5 minutes after which the supernatant was diluted 1:5 in binding buffer (20 mM sodium phosphate, pH 7.0 – 7.4) and filtered (0.45 µm). Samples were passed through a HiTrap protein G HP column (GE healthcare, Sweden) following manufacturer instructions. After washing with binding buffer, total IgGs were eluted (0.1 M glycine HCl, pH 2.7) and collected in neutralisation buffer (1M Tris, pH 9). The buffer of the purified IgGs was exchanged to PBS overnight at 4°C using Spectra/Por 6 dialysis membrane (VWR), centrifuged at 3000 g (5 min, 4°C) and filtered over 0.22 µm Milex low protein binding durapore filters (Millipore). The included patients tested also positive for anti-FHL1 antibodies.

***Force and $[Ca^{2+}]_i$ measurements in single fibers***

In some experiments, FDB muscles treated with serum as above were mechanically dissected into single fibres to enable measurements of contractile properties and intracellular $Ca^{2+}$ ($[Ca^{2+}]_i$). The tendons were attached to aluminium T-clips and mounted to hooks connected to an Aurora 403A force transducer in a stimulation chamber. The single muscle fibres were continuously superfused with bubbled (95% $O_2$ and 5% $CO_2$ to give a pH of 7.3) Tyrode buffer solution (room temperature) when mounted in the stimulation chamber. The fibres were stretched until optimal length was reached and then stimulated according to a force protocol. To measure $[Ca^{2+}]_i$, the fluorescent indicator Indo-1 AM was used. Indo-1 was excited at 360 nm and emitted at 405 and 495 nm wavelength and these signals were translated into into $[Ca^{2+}]$ using a calibration method[3 4]. Fiber cross-sectional area (CSA) was calculated by measuring fiber diameter using a camera mounted on the microscope. Specific force and $[Ca^{2+}]$ was measured simultaneously by stimulating the fibers at increasing frequencies from 15-100 Hz at 350 ms duration. Serum from 3 patients and 2 healthy controls were used for these experiments.
